# Isoform-level gene expression patterns in single-cell RNA-sequencing data

**DOI:** 10.1101/036988

**Authors:** Trung Nghia Vu, Quin F Wills, Krishna R Kalari, Nifang Niu, Liewei Wang, Yudi Pawitan, Mattias Rantalainen

## Abstract

RNA-sequencing of single-cells enables characterization of transcriptional heterogeneity in seemingly homogenous cell populations. In this study we propose and apply a novel method, ISOform-Patterns (ISOP), based on mixture modeling, to characterize the expression patterns of pairs of isoforms from the same gene in single-cell isoform-level expression data. We define six principal patterns of isoform expression relationships and introduce the concept of differential pattern analysis. We applied ISOP for analysis of single-cell RNA-sequencing data from a breast cancer cell line, with replication in two independent datasets. In the primary dataset we detected and assigned pattern type of 16562 isoform-pairs from 4929 genes. Our results showed that 78% of the isoform pairs displayed a mutually exclusive expression pattern, 14% of the isoform pairs displayed bimodal isoform preference and 8% isoform pairs displayed isoform preference. 26% of the isoform-pair patterns were significant, while remaining isoform-pair patterns can be understood as effects of transcriptional bursting, drop-out and biological heterogeneity. 32% of genes discovered through differential pattern analysis were novel and not detected by differential expression analysis. ISOP provides a novel approach for characterization of isoform-level expression in single-cell populations. Our results reveal a common occurrence of isoform-level preference, commitment and heterogeneity in single-cell populations.

## INTRODUCTION

The emergence of single-cell RNA sequencing (scRNAseq) enables characterization of gene expression variability on the single-cell level (1, 2). Prior to the advent of scRNAseq, typical gene expression measurements were only possible based on the average expression level over a large number of cells (bulk-cell RNAseq), which effectively excluded the possibility to study gene expression heterogeneity at the single cell level.

Single-cell sequencing has been applied in a wide range of research areas to date, including studies of circulating tumor cells (1, 3), breast cancer (4), prostate cancer (5), transcriptional dynamics (6), cell cycle (7), tissue heterogeneity (8) and cell-to-cell variation in alternative splicing via isoform-level expression analysis (9–11). Multiple recently published reviews (2, 12, 13) provide excellent and broad introduction to single-cell sequencing.

Transcriptional isoforms are defined as mRNA molecules of different length and exon composition originating from the same locus, which code for multiple forms of the corresponding protein. Transcriptional isoforms arise as mRNAs are produced from different transcriptional starting sites, terminated at different polyadenylation sites or as a consequence of alternative splicing (14, 15). There are numerous studies of alternative splicing in the context of bulk-cell RNAseq, including studies of tissue-level regulation of isoform expression (16) and prediction and quantification of alternative isoforms (17, 18).

To date there are relatively few studies published that are focused on characterization of isoform-level expression at the single-cell level. Potentially novel splice junctions were discovered after studying alternative splicing in single cells (10, 19). Shalek et al. (9) described bimodality in the expression of genes and isoforms in scRNAseq data and preference of individual cells to express a particular isoform from genes with multiple isoforms was investigated, although this study was based on a limited dataset with RNAseq data from only 18 cells. In another study (11), statistical modeling was applied to characterize 3’ isoform choice variability in single cells via a transcriptome-wide analysis of scRNAseq data from 48 single-cells using BATSeq (11), which is a sequencing methodology with a prominent 3’ end sequencing bias. Recently, Welch et al (20) introduced a statistical model to detect isoform usage that shows significant biological variation through the contrast of variance of isoform ratios to technical noise estimated from alternative splicing modules.

In this study we propose a novel method, ISOform-Patterns (ISOP), for analysis and characterization of single-cell isoform-level gene expression data. ISOP enables analysis of single-cell preference, commitment and heterogeneity of isoform level expression. Based on this method, we defined a set of six principal patterns of isoform expression relationships between isoforms from the same gene, including isoform preference, bimodal isoform preference, and mutually exclusive expression commitment. We apply ISOP for analysis of scRNAseq data from a breast cancer cell line (MDA-MB-231, N=327 cells), with replication in two independent single-cell datasets, with the aim of systematically characterizing the extent and nature of single-cell isoform-level expression patterns.

We also assess to what extent isoform patterns arise randomly due to the distributional properties of single-cell RNA expression, and finally we demonstrate how ISOP can be applied for differential isoform pattern (DP) analysis.

## MATERIAL AND METHODS

### Datasets

The primary dataset include 384 single-cell RNA sequencing samples from triple-negative breast cancer cell line (MDA-MB-231) of which half were treated with metformin. Specifically, MDA-MB-231 cell was cultured in ATCC-formulated Leibovitz’s L-15 Medium (Manassas, VA) supplemented with 10% fetal bovine serum (FBS, Atlanta Biologicals, Flowery Branch, GA) and incubated at 37°C without CO_2_. Cells were plated into 6-well plate at a seeding density of 6 x 10^4^ cells/well and were treated with or without 1mmol/L metformin (Sigma-Aldrich, St. Louis, MO) after 24 hours of incubation. Fresh medium and drug were replaced every 24 hours. After 5 days of drug treatment, cells were resuspended and single-cells were captured using the Fluidigm C1 system immediately. Two independent cell culture batches were used from which 2 x 96 untreated cells (control) were captured, and 2 x 96 treated cells were captured. Furthermore, cells were captured on two different C1 machines in an orthogonal design in relation to the treatment groups. Sequencing libraries were prepared using the standard Fluidigm protocol based on SMARTer chemistry and Illumina Nextera XT. RNA sequencing of 100 bp paired-end reads was carried out on an Illumina HiSeq with 4.9 million reads / cell on average.

The first public dataset consists of 96 cells from HTC116 cell-line extracted from a public dataset (21). Single-cells were captured using the Fluidigm C1 system and sequencing libraries for Illumina sequencing were prepared based on SMARTer chemistry and Illumina Nextera XT. The 96 libraries, divided into two pooled samples of 48 libraries and sequenced on two lanes on a Illumina HiSeq. For further details, see the original paper (21).

The second public dataset include 305 cells from primary human myoblasts (6) after eliminating samples with debris, without cells and or many cells (bulk-cell). Single-cells were captured using the Fluidigm C1 system and sequencing libraries for Illumina sequencing were prepared based on SMARTer chemistry and Illumina Nextera XT. The further details, see the original publication (6).

### Data preprocessing

The Fastq files from the primary dataset (MDA-MB-231) for single-cell RNAseq were processed through MAP-RSeq pipeline (22) to assess the quality of reads which includes determination of cells with no or low reads, assessment of duplicate reads, inspection of gene-body coverage, estimation of distance between paired-end reads, and evaluation of sequencing depth. The Fastq files were mapped to human hg19 UCSC annotation reference using Tophat (23) and Bowtie (24) to create bam files. In all further analyses, we used the same type of annotation reference but updated from igenomes (http://support.illumina.com/sequencing/sequencingsoftware/igenome.html). In practice, only annotation information of chromosomes chr1 to chr22, chrX and chrY were used to extract isoform and gene expression. Cufflinks (18) version 2.2.1 were used to estimate the abundances of gene and isoform expression from the bam files. For the public datasets, we also applied Tophat (23) and Bowtie (24) with the same annotation reference for read alignments, and further Cufflinks (18) for quantifying isoform level expression values.

The MDA-MB-231 dataset was subsequently preprocessed further by excluding 57 samples corresponding to empty wells (39 samples) and atypical samples (18 outliers) identified by Principal Component Analysis. In addition, the isoforms expressed in less than 1% of samples were filtered out from the further analysis. Isoforms were considered as expressed if their read counts were ≥ 3 in a cell. After pre-processing the isoform-level dataset contained 21728 isoforms, of which 13863 isoforms from 4929 genes were multiple-isoform genes, and scRNAseq data from 327 single-cells were available for analysis, of which 162, nearly a half of samples, were from the metformin treated group. Gene-level gene expression of this dataset comprised 13073 genes that were also extracted from the Cufflinks. The HTC116 dataset consisted of 26723 isoforms from 96 single cells and the myoblast dataset included 25623 isoforms from 305 single cells.

### Patterns of isoform expression

To characterize isoform-level gene-expression patterns in scRNAseq data and to detect potential subpopulations of cells, log expression differences Δ_*i,j,a,b*_ (Eq 1) between pairs of isoforms {*a*, *b*} of a single gene (*j*) at cell *i* from a population of *N* cells (*i* = 1 … *N*) were modeled using a Gaussian mixture model approach (Eq 2-3). Where *y_i,j,a_* and *y_i,j,b_* represent the log expression of isoform *a* and isoform *b* in cell *i*, parameter *w_k_* is the mixing weight for component *k* in the model and *K* is the total number of components in the model. In our analyses *K* was constrained to ≤ 3. For simplicity, indexes relating to gene (*j*) and cell (*i*) were omitted from Eq 2 and Eq 3.

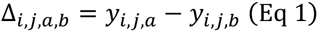

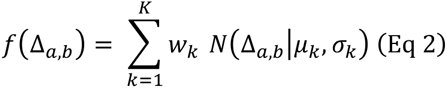

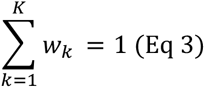

Model selection to determine the number of components of each mixture model was done by Akaike Information Criterion (AIC) scores with the additional requirement that the smallest weight (w_k_) had to be >0.025, and that the standard deviation of all components were greater than 0.01. Mixture models were fitted using a computationally efficient histogram-based method implemented in the OCplus package (25). Fitting of the mixture models using the OCplus algorithm reduces data to a histogram defined by equally spaced bins weighted by the number of data points in each bin, here the number of bins was set to the square root of the number of data points (number of cells). Based on the mixture model approach, we define six principal isoform-pair patterns (see **Results** section). The method, ISOform-Patterns (ISOP), was implemented in the R package *ISOP* (version 0.99.0 was used in the analyses in this study), available at (https://github.com/nghiavtr/ISOP) under a GPL-3 license.

### Test for non-randomness of isoform-pair distributions

To test if an isoform pair distribution was significantly non-random, we applied the χ^2^ goodness-of-fit test combined with a permutation-based approach. For an isoform pair {*a*, *b*}, we permute the isoform vectors and calculate Δ_*a,b,perm*_, which is the expression difference between the permuted isoform *a* and isoform *b* vectors, 10000 permutations were applied. Next, we estimate the mean, *E*(Δ_*a,b,perm*_), from the permutations and for each bin. The permutation-based null distribution was derived from the χ^2^ goodness-of-fit test of *k*th permuted isoform pairs Δ_*a,b,perm*(*k*)_ and *E*(Δ_*a,b,perm*_). The observed test statistic was derived from the χ^2^ goodness-of-fit test between Δ_*a,b*_ and *E*_*a,b,perm*_). Finally, a p-value was computed by comparing the observed test statistic with the permutation-based null distribution of the χ^2^ statistic. To determine significant (non-random) isoform-pairs, false discovery rate (FDR) was estimated from p-values using Benjamini & Hochberg (BH) multiple testing method (26).

### Differential pattern analysis

We test if a treatment effect (metformin exposure) was associated with the probability of cells of being clustered into a particular mixture model component in the isoform pattern models, which would suggest a treatment effect on the distribution of isoform pairs. For each mixture model with more than one component, we assigned individual cells to components (cluster labels) based on the estimate of the probability that cell *i* belongs to component *k*. Subsequently, we applied a permutation test to test the association between the cluster labels and the metformin treatment status. In the permutation test we permuted the metformin treatment factor (10000 permutations). Next, we established a null-distribution from the χ^2^ statistics of chi-squared test between the cluster labels and the permuted group labels. Finally, we computed the χ^2^ statistic of chi-squared test between cluster labels and the true group labels, and compared it to the permutation-based null distribution to obtain a permutation-based p-value. False discovery rate (FDR) was estimated from the p-values using Benjamini & Hochberg (BH) multiple testing method (26).

### Differential expression analysis

To identify differentially expressed (DE) isoforms, we applied a linear model as implemented in the limma package version 3.22.1 (27), accounting for the mean-variance trend in the data, which has also be applied for analysis of RNAseq data (28). Prior to the differential expression analysis, the count dataset was expressed as log-counts per million (cpm) using functionality in the edgeR package version 3.8.2 (29). We accounted for culture batch effect and machine effect in the experimental design matrix in the limma analysis of the primary dataset. We applied a permutation test to identify DE isoforms. First, 100 permuted datasets were generated by randomly permuting group labels. Next, the moderated t-statistics (extracted from limma) of isoforms from the actual dataset and the population of the moderated t-statistics of the isoforms from permutated datasets were used to calculate empirical p-values of the isoforms. Finally, false discovery rate (FDR) were estimated by adjusting empirical p-values using Benjamini & Hochberg (BH) correction (26). DE isoforms were defined as isoforms with FDR values less than or equal to 5%.

## RESULTS

We developed and applied a method (ISOP) for transcriptome-wide analysis of the co-variability of expression levels in pairs of isoforms (*a* and *b*) from the same gene in scRNAseq data. ISOP utilizes a Gaussian mixture model approach to model isoform expression difference (Δ_*a,b*_) on a log scale (see **Methods** for details). Based on the estimated mixture model parameters including the number of components and the location of the components, expression patterns of pairs of isoforms in individual genes can be systematically characterized and described by a small set of principal isoform expression patterns. The isoform expression patterns can be interpreted in terms of single cell isoform expression preference, commitment and heterogeneity.

### Principal patterns

Based on ISOP, we define six distinct patterns of isoform expression (Figure 1) with characteristics outline in the rest of this section.

**Figure 1.**
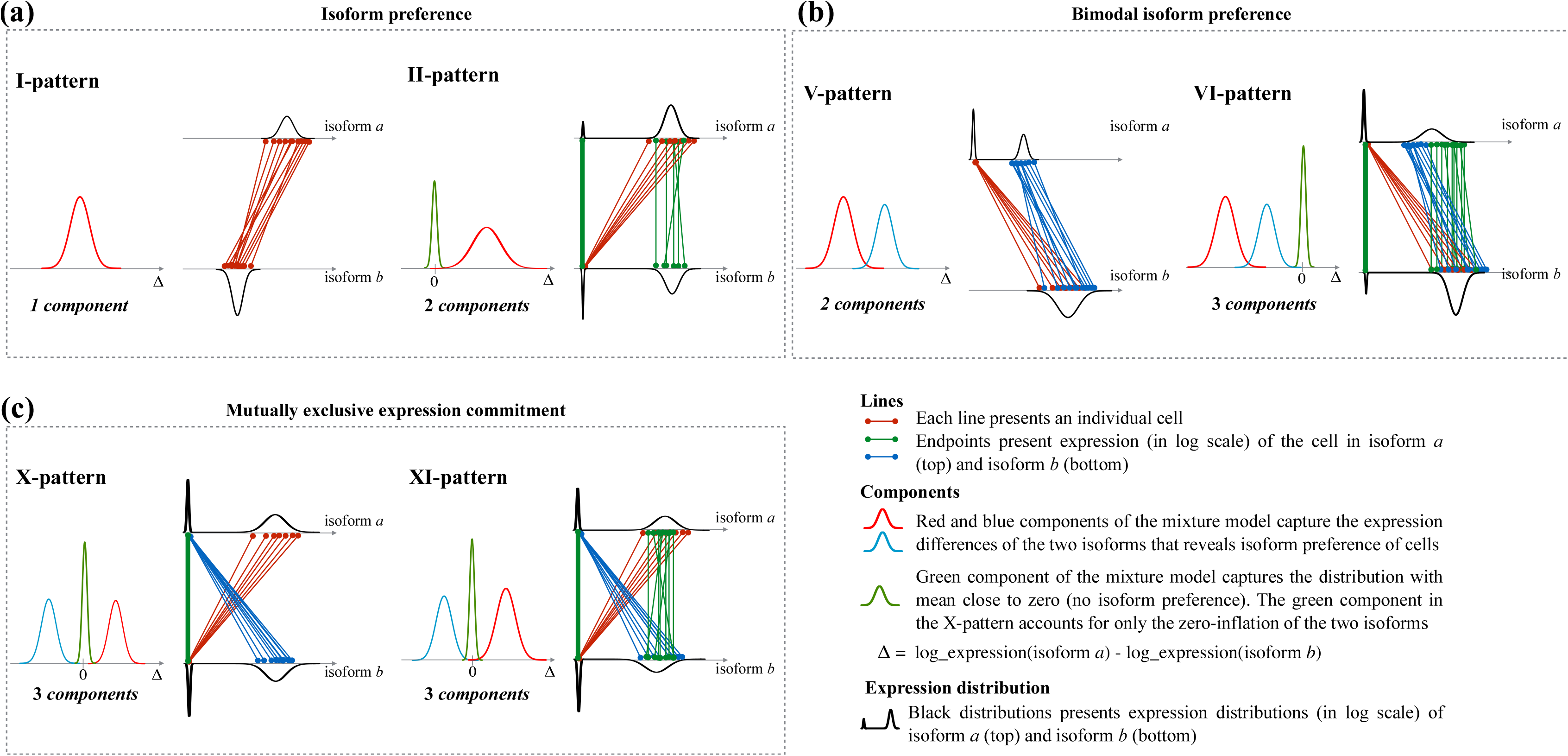
Overview of the six principal isoform expression pattern types. Each panel consists of two plots: a component plot (left) displaying the typical mixture model of Δ_*a,b*_) for the pattern, corresponding to isoform *a* and isoform *b* in the isoform pair, and a pair-line plot (right) of the two isoforms. (a) The I-pattern (isoform preference of cells) and its extension, the II-pattern (isoform preference in a subset of cells). (b) The V-pattern (bimodal isoform preference of cells) and its extension, the VI-pattern (bimodal isoform preference in a subset of cells). (c) The X-pattern (mutually exclusive expression commitment of cells) and its extension, the VI-pattern (mutually exclusive expression commitment in a subset of cells).

#### I-pattern

A single-component model defines this pattern. Thus, there is no cell-to-cell heterogeneity in the isoform pair. However, if the mean of the mixture component is distant from zero, this indicates that one isoform is preferred over the other isoform *(preference)*.

#### II-pattern

This pattern is an extension of the I-pattern with an additional mixture component capturing the zero-inflation of cells where isoform expression is not detected, or where isoforms are expressed at close to equal amounts in both isoforms. The II-patterns represents isoform preference in a subset of cells.

#### V-pattern

A two-component mixture model defines the V-pattern, in which the means of the two components share the same sign. However, unlike the I-pattern, which has a unimodal distribution in both isoforms, the V-pattern generally has a unimodal expression in one of the isoforms, while the other isoform bimodal distribution. Thus, the V-pattern is defined by a bimodal isoform preference that indicates cell-to-cell heterogeneity caused by prominent bimodality in one of the two isoforms.

#### VI-pattern

This pattern is an extension of the V-pattern with an additional component in the mixture model with its mean close to zero (in analogy to how the I-pattern extends the II-pattern). Thus, the VI-pattern represents the bimodal isoform preference of a subset of cells in population.

#### X-pattern

A three-component mixture model defines the X-pattern. Different from the previous pattern types, the X-pattern has two components with the location parameters (mean) of opposite sign and a third component accounting for the zero-inflation. This pattern captures pairs of isoforms with mutually exclusive expression (mutually exclusive isoforms, MXIs), with similarities to the concept of mutually exclusive exons (MXEs) (16). The X-pattern represents a mutually exclusive expression commitment that can be interpreted as an indication of commitment of individual cells to express either one of the isoforms, but not both, representing a particular type of inter-cell heterogeneity.

#### XI-pattern

This is an extension to the X-pattern where the component located close to zero accounts for both zero-inflation and cells where the two isoforms are expressed at close to equal. Thus, the XI-pattern represents a mutually exclusive expression commitment in a subset of the cell population.

### Classification and observed frequencies of isoform patterns

We applied ISOP for analysis of 13863 isoforms from 4929 multiple-isoform genes in the MDA-MB-231 single-cell dataset. We detected and assigned pattern type to 16562 isoform-pairs (Figure 2a). We found that 0.2% of the isoform pairs were classified as I-pattern and 8.1% pairs were classified as the related II-pattern. More than a half (55.2%) of the I-patterns were consistent with isoform preference, defined by the absolute mean of the mixture component >0.5 on the log scale, marked by stars in the panel I-pattern of Figure 3a. In contrast, the V-pattern and its extension, the VI-pattern, had proportions in a similar range, 5.7% and 8.2%, respectively (13.9% in total). The X-pattern and XI-pattern were the most common patterns, accounting for 77.9% of the isoform pairs, of which 17.2% were MXIs.

**Figure 2.**
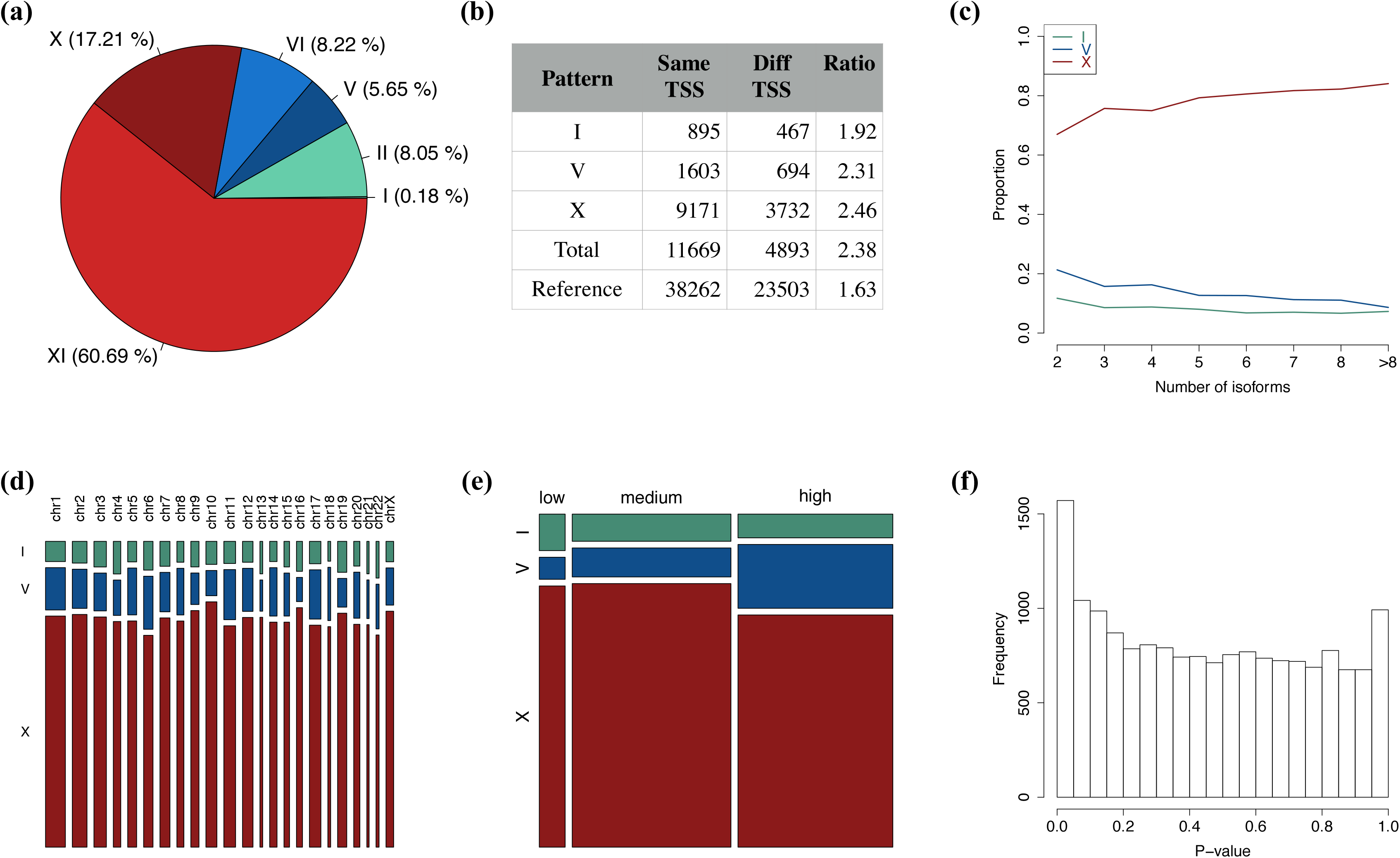
ISOP analysis of the MDA-MB-231 dataset. (a) Proportion of isoform patterns. (b) Frequency of patterns with isoforms with the same and different transcription start site (TSS). (c) Proportion of patterns as a function of the total number of annotated isoforms in the corresponding gene. (d) Proportion of patterns stratified by chromosome. (e) Proportion of patterns stratified by gene expression level. (f) P-value distribution from the test of association between component label and treatment group.

**Figure 3.**
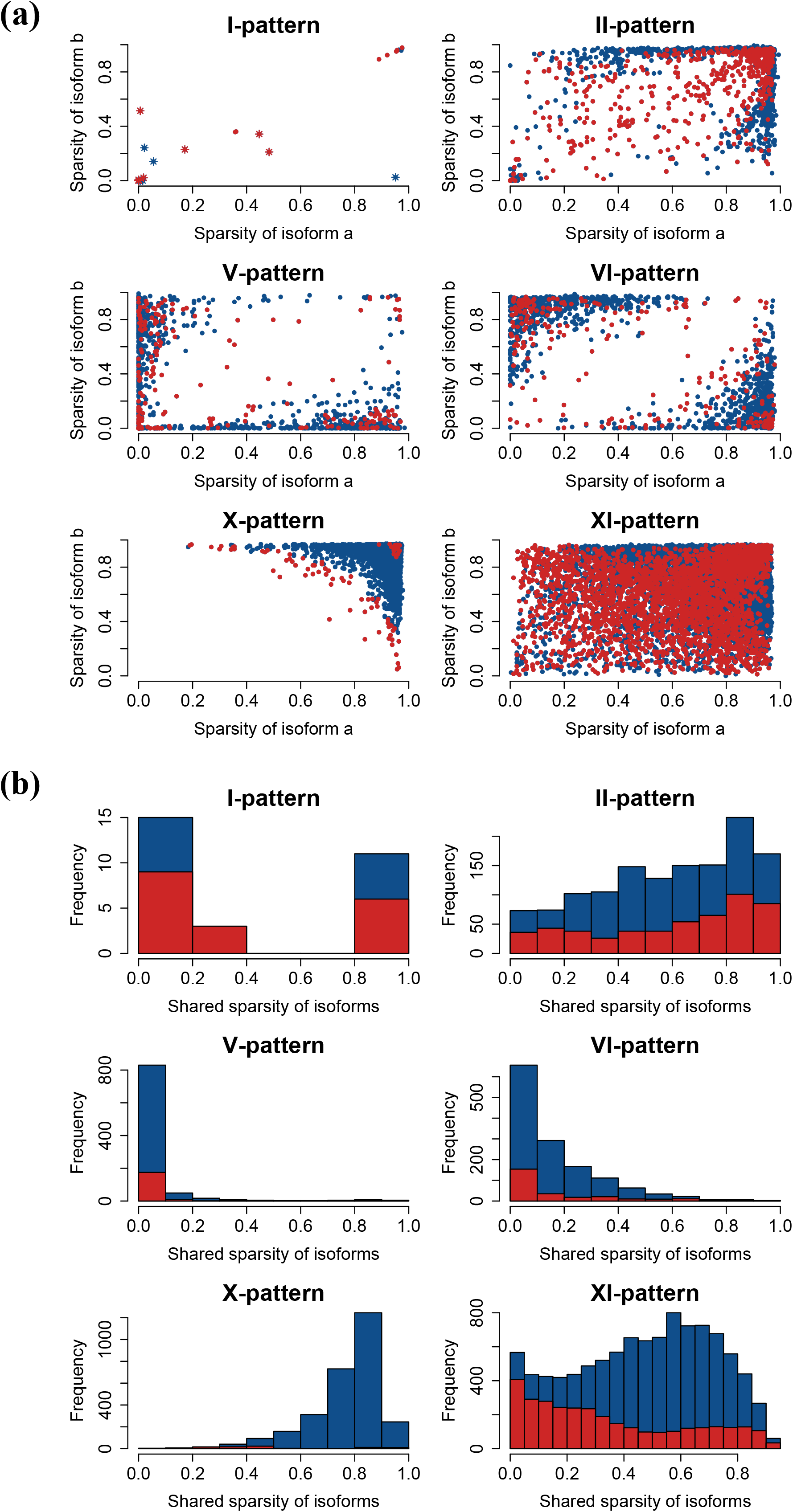
Sparsity of isoform expression in principal isoform expression pattern types. (a) Pairwise sparsity of isoform *a* and isoform *b* in each individual pair and pattern type. Each point presents a single isoform-pair pattern, while the sparsity of the two isoforms in the pair are indicated on the x-axis and the y-axis. Blue points and red points represent non-significant and significant isoform-pairs. The star points indicate I-patterns with isoform preference. (b) Empirical distribution of the shared sparsity of the two isoforms (directly related to the sparsity of Δ_*a,b*_) in isoform pairs across pattern types. The red part of the histograms corresponds to the portion of significant (non-random) isoform-pairs.

We applied a permutation test (see **Methods**) to assess to what extent isoform-pair patterns were significant (non-random, FDR ≤ 0.05). We found 4309 (26.0%) significant isoform patterns in total (Table 1). The I-pattern and II-pattern were the least common patterns (Figure 2a), while they had the highest proportions of significant isoform-pair patterns, 62.1% and 39.3% for I-pattern and II-pattern respectively. The X-patterns were the second most common pattern (Figure 2a), while only 3.1% of these were statistically different from the permutation-based null distribution. A total of 3213 (32.0%) of the XI-pattern isoform-pairs were found to be significant, representing the most commonly observed significant isoform pattern.

**Table 1.**
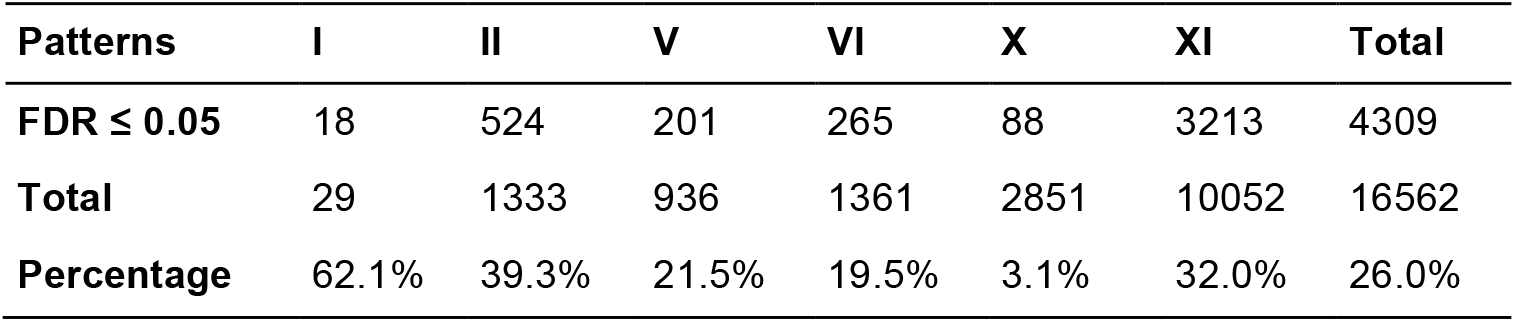
Patterns with significant (non-random) isoform-pairs (FDR ≤ 0.05) in the MDA-MB-231 dataset.

| Patterns | I | II | V | VI | X | XI | Total |
| --- | --- | --- | --- | --- | --- | --- | --- |
| **FDR ≤ 0.05** | 18 | 524 | 201 | 265 | 88 | 3213 | 4309 |
| **Total** | 29 | 1333 | 936 | 1361 | 2851 | 10052 | 16562 |
| **Percentage** | 62.1% | 39.3% | 21.5% | 19.5% | 3.1% | 32.0% | 26.0% |

Next, we investigated how common expression sparsity induced isoform patterns were. We define sparsity of expression as the proportion of cells where expression levels were below detection limit, so that an isoform with low sparsity has detectable expression levels from most cells. We found that the X-pattern was mainly detected in isoform pairs with high sparsity (Figure 3a), suggesting that the great majority of X-pattern isoform pairs are likely to arise as a direct consequence of sparsity, which can be caused by e.g. transcriptional bursting, biological cell-to-cell heterogeneity or due to the sensitivity limitations of scRNAseq (including transcript drop-out effects). Next, we assessed the distribution of the proportion of cells with zero detected reads in both isoforms in a pair, a quantity directly related to the sparsity of Δ_*a,b*_, which is the quantity modeled in the mixture models (Figure 3b). It is evident that the significant patterns (in red color) usually consist of two low sparsity isoforms, with the exception of the I-pattern and II-pattern. Thus, overall 26.0% of all isoform-pairs were found to be significant (non-random), while a large fraction of non-significant isoform-pairs are likely to be induced by sparsity in the isoform-level expression data.

To replicate these results, we applied the ISOP method for analysis of two additional public datasets (HTC116 dataset and myoblast dataset, see **Methods**) and found that the same patterns were also present in these data sets. There were 23712 and 22763 isoform pairs in the HTC116 dataset and myoblast dataset respectively. The proportions of patterns in these datasets (Figure 4a and Figure 4b) were similar to the proportions observed in the MDA-MB-231 dataset, where the X-pattern and the XI-pattern were the most common. The HTC116 dataset and the myoblasts dataset (Table 2) had 5.1% and 22.8% significant patterns, respectively. Similar to the results of the MDA-MB-231 dataset, the non-random isoform-pairs that were annotated as X-patterns in the both datasets account for the smallest proportions as compared to other patterns, 0.6% in the HTC116 dataset and 3.0% in the myoblast dataset.

**Figure 4.**
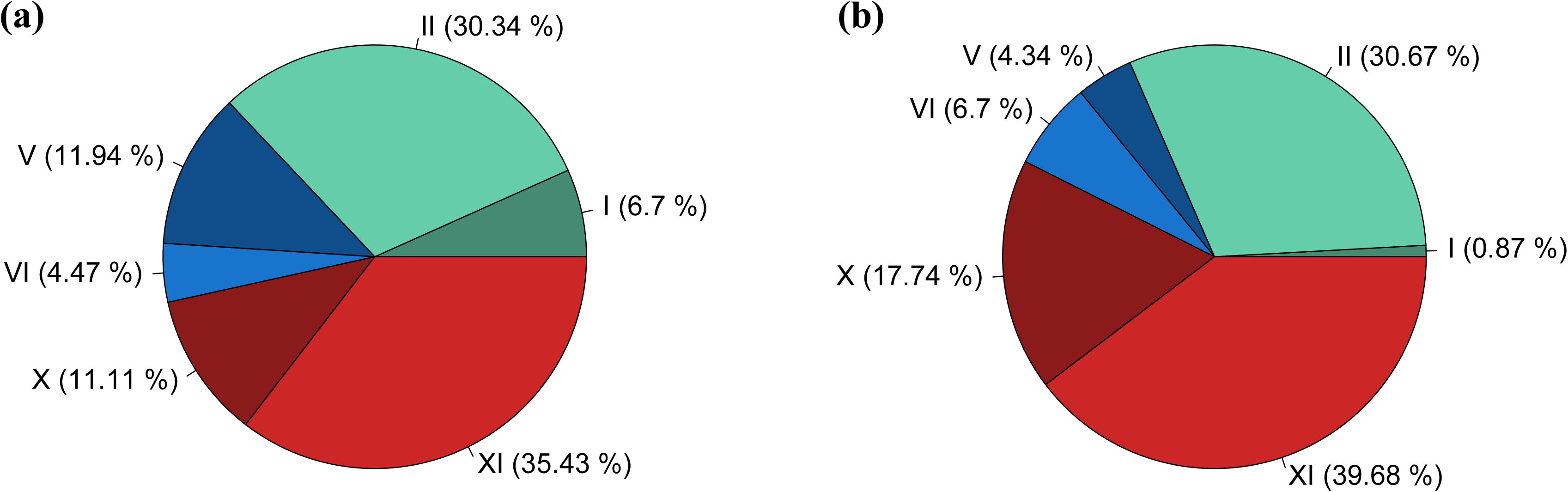
Proportion of patterns detected in replication data sets. (a) HTC116 dataset. (b) myoblast dataset.

**Table 2.** Patterns with significant (non-random) isoform-pairs (FDR ≤ 0.05) in the public datasets.

|  | Patterns | I | II | V | VI | X | XI | Total |
| --- | --- | --- | --- | --- | --- | --- | --- | --- |
| HTC116 | **$FDR \leq 0.05$** | 44 | 312 | 110 | 26 | 17 | 704 | 1213 |
|  | **Total** | 1589 | 7195 | 2832 | 1059 | 2635 | 8402 | 23712 |
|  | **Percentage** | 2.8% | 4.3% | 3.9% | 2.5% | 0.6% | 8.4% | 5.1% |
| Myoblast | **$FDR \leq 0.05$** | 76 | 1239 | 214 | 253 | 122 | 3294 | 5198 |
|  | **Total** | 197 | 6981 | 989 | 1525 | 4039 | 9032 | 22763 |
|  | **Percentage** | 38.6% | 17.7% | 21.6% | 16.6% | 3.0% | 36.5% | 22.8% |

### Association of isoform patterns with genomic features

We then assessed if isoform patterns were associated with different transcription start site (TSS), the number of annotated isoforms of the gene, chromosome location and average expression level. In following analyses we merge I, V and X patterns with their corresponding extension patterns. We observed that the number of isoform pairs originating from the same TSS is more than twice as common (2.38 times, across all patterns) compared with pairs originating from different TSS, compared to a ratio of 1.63 times observed in the reference transcriptome (Figure 2b). Thus, across all detected patterns, isoform pairs from the same TSS are more frequently observed than those originating from different TSS. The number of isoforms in each gene positively correlates with the proportion of the X-pattern (r = 0.92), negatively correlates with the proportion of I-pattern and V-pattern (r = -0.82 and r = -0.94) (Figure 2c). Hence, genes with many isoforms tend to have a higher proportion X-patterns and lower proportion I-pattern and V-pattern. There was no association between pattern type and the chromosome on which the gene is located (Figure 2d). Finally, we investigated to what degree the different isoform patterns were associated with gene-level expression. We grouped genes into three groups: low, medium and high expression defined by quartiles of average gene expression (<1^st^ quartile, >1^th^ quartile & <3^rd^ quartile, >3^rd^ quartile). Most isoform pairs belonged to the high expression group (45.6%) or the medium expression group (46.8%); while only a small proportion (7.6%) of these are from low expression group (Figure 2e).

### Isoform patterns provide a novel way to assess biological effects in scRNAseq data

Next, we tested for associations between a treatment (metformin exposure) and proportion of cells in each mixture model component (see **Methods** for details). An association between a treatment effect and the proportion of the cells in each mixture component would indicate an effect of the treatment on the isoform-level expression pattern. Figure 2f displays the distribution of empirical p-values from the association tests, suggesting an enrichment of low p-values. A total of 80 isoform pairs were defined as significant (FDR ≤ 0.05) from 54 genes. We define these genes as differential-pattern genes (DP genes) to distinguish them from differentially expressed genes (DE genes). Of these isoform pairs 19 (23.8%) were found to be significant and non-random (FDR ≤ 0.05) in respect to the distribution of Δ_*a,b*_, using the previously described permutation test. However, non-significant results in respect to the distribution of Δ_*a,b*_ does not provide evidence that excludes the possibility of true treatment effects on the proportion of treated cells in the mixture components in these patterns, only that the overall distribution of Δ_*a,b*_, could have occurred by chance or induced by e.g. sparsity. A total of 37 (68.5%) of the DP genes had at least one of the two isoforms differently expressed (DE), and 5 of these DP genes were from isoform pairs (9.3%) with two DE isoforms. Furthermore, 17 DP genes (31.5%) did not have either of the two isoforms differentially expressed. The DP unique genes were: *NABP1, SMN2, BTN3A3, CLIC1, CEP85L, ERLIN2, CDK1, TMEM136, DAZAP2, PMP22, SPECC1, ACTG1, NFIC, TNPO2, NFATC2, CDC45*, and *BCAP31*.

## DISCUSSION

A unique property of single-cell transcriptomic profiling is the ability to characterize cell-to-cell heterogeneity in cell populations. Our objective was to investigate cellular heterogeneity in isoform-level gene expression based on scRNAseq profiling. We proposed a novel method, ISOform-Patterns (ISOP), using a mixture model to model and categorize isoform pairs into principal isoform expression patterns.

We described six principal patterns of isoform expression, which interpretation in respect to isoform preference, bimodal isoform preference and mutually exclusive isoform expression commitment. Each pattern type represents a specific expression relationship between a pair of isoforms from the same gene. The I-pattern characterizes isoform preference in the cell population of one isoform over the other isoform. The V-pattern expresses a bimodal isoform preference indicating cell-to-cell heterogeneity associate with one level of expression of one isoform and two levels of expression of the other isoform in the cell population. The X-pattern describes a mutually exclusive expression commitment pattern of the cells to express either one of the isoforms, but not both. The II-pattern, VI-pattern and XI-pattern are extensions of I-pattern, V-pattern and X-pattern respectively where a subset of cells display the pattern. The type of isoform preference of cells reported in previous studies (9, 11) can be accounted for by the I-pattern, V-pattern, or their respective extensions. Isoform commitment, as defined by mutually exclusive isoform expression (the X-pattern and XI-pattern) was the most common patterns observed, assigned to 77.9% of the isoform pairs. We showed that a large proportion (26.0%) of isoform pair patterns were found to be statistically significant (non-random), while remaining patterns (74.0%) might have been stochastically generated, mainly as a function of the sparsity (zero inflation) or the degree of bimodality in the isoform expression distribution. Such sparsity can arise due to transcriptional bursting, biological heterogeneity or transcript drop-out effects or other technical limitations inherit to scRNAseq.

We also outlined how the ISOP method can be applied to test for biological effects related to the principal isoform expression patterns, which was represented by a small molecule perturbation effect (metformin exposure) in the primary dataset. DP analysis provides a novel approach to detect isoform-related effects that may not have been discovered through conventional DE analysis. We discovered 54 significant DP genes, of which 31.5% were associated with isoforms that were not differentially expressed. Thus, significant DP genes constitute novel information that augments traditional analyses.

Our study has some limitations. Firstly, the analysis is focused on the set of annotated isoforms only and we indirectly assume that annotations are correct. Secondly, the present study did not have ERCC spike-ins that could be used to establish levels of technical noise in the data. Furthermore, many algorithms have been proposed for quantification of isoform level gene expression from RNAseq data (18, 30, 31), and are all based on slightly different assumptions. In our analyses we applied the widely used Cufflink software for isoform expression estimation. Isoform-level gene expression quantification is inherently more challenging than gene level quantification, particularly in scRNAseq analysis where there are limited number of RNAseq reads from each cell, and one would expect a degree of variability associated with the quantification algorithm applied. Furthermore, we make the assumption that the expression differences between pairs of isoforms on a log scale can be approximated by a Gaussian mixture model. Finally, in this study we have focused on modeling pairwise isoforms expression patterns, while commitment of cells in set with more than two isoforms is also interesting and biologically relevant, something that we are interested in exploring in future studies.

In conclusion, ISOP provides a novel approach for characterizing isoform-level expression in single-cell populations. ISOP also introduces a novel approach to discover DP genes associated with biological effects, which is complementary to conventional analysis of differential expression.

Although isoform expression patterns can arise as a function of sparseness in expression patterns, we found that more than a quarter of the patterns in our dataset were found to be non-random, suggesting common occurrence of isoform-level preference, commitment and heterogeneity in singlecell populations.

## ACKNOWLEDGEMENT

This work was supported by grants from the Swedish Research Council (Unga Forskare) and Swedish e-Science Research Centre (SERC) – “e-Science for Cancer Prevention and Control (ecpc)” to MR; and grants from the Swedish Research Council and the Swedish Cancer Foundation to YP.

